# Intellectual Disability-related genes increase ADHD risk and locomotor activity in *Drosophila*

**DOI:** 10.1101/725937

**Authors:** Marieke Klein, Euginia Singgih, Anne van Rens, Ditte Demontis, Anders D. Børglum, Nina Roth Mota, Anna Castells-Nobau, Lambertus A. Kiemeney, Han G. Brunner, Alejandro Arias-Vasquez, Annette Schenck, Monique van der Voet, Barbara Franke

## Abstract

**Objective:** Attention-Deficit/Hyperactivity Disorder (ADHD) is a common, highly heritable neuropsychiatric disorder. ADHD often co-occurs with Intellectual Disability (ID), and shared overlapping genetics have been suggested. This study aimed to identify novel ADHD genes by investigating whether genes carrying rare mutations linked to ID contribute to ADHD risk through common genetic variants. Validation and characterization of candidates were performed using *Drosophila melanogaster*.

**Method:** Common genetic variants in a diagnostic gene panel of 396 autosomal ID genes were tested for association with ADHD risk, through gene-set and gene-wide analyses, using ADHD meta-analytic data of the Psychiatric Genomics Consortium (n=19,210) for discovery and iPSYCH ADHD data for replication (n=37,076). The significant genes were functionally validated and characterized in *Drosophila* by assessing locomotor activity and sleep upon knockdown of those genes in brain circuits.

**Results:** The ID gene-set was significantly associated with ADHD risk in the discovery and replication data-sets. The three genes most consistently associated were *MEF2C*, *ST3GAL3*, and *TRAPPC9*. Performing functional characterization of the two evolutionary conserved genes in *Drosophila melanogaster*, we found their knockdown in dopaminergic (*dMEF2*) and circadian neurons (*dTRAPPC9*) to result in increased locomotor activity and reduced sleep, concordant with the human phenotype.

**Conclusions:** This study reveals that a large set of ID-related genes contributes to ADHD risk through effects of common alleles. Utilizing this continuity, we identified *TRAPPC9, MEF2C*, and *ST3GAL3* as novel ADHD candidate genes. Characterization in *Drosophila* suggests that *TRAPPC9* and *MEF2C* contribute to ADHD-related behavior through distinct neural substrates.

## INTRODUCTION

Attention-Deficit/Hyperactivity Disorder (ADHD) is a common neurodevelopmental disorder with prevalence estimates of 5.3% in childhood and 2.5–4.9% in adulthood(1). ADHD is clinically characterized by two core symptom domains: inattention and hyperactivity/impulsivity, which can occur individually or combined(1). Despite the high heritability (70–80%)(1), identification of ADHD risk genes has been difficult, mainly due to ADHD’s complex genetic architecture(1). Genetic variants that occur frequently in the population and have generally small individual effects on disease risk are thought to underlie the disorder in most patients, and first genome-wide significant findings for ADHD have been identified only recently(2).

Intellectual Disability (ID) refers to a highly heterogeneous group of childhood-onset disorders characterized by below-average intellectual functioning (IQ<70) and significant limitations in adaptive functioning, which covers many everyday social and practical skills(3). ID has an estimated prevalence of 2–3% in the population; severe handicaps have a population-prevalence of 0.3–0.5%(4). ID is often monogenic, but many different genes and types of mutations are implicated(3). ADHD is a common comorbid disorder in children with ID(5). Studies of children with mild and borderline ID have identified ADHD in 8–39% of the cases(5). A recent study using the Swedish birth registry data showed that nearly all of this comorbidity can be attributed to genetic factors(6). Based on such phenotypic and genetic overlap, it has been hypothesized that ID and ADHD, and neurodevelopmental disorders more broadly, have an overlapping genetic etiology(6).

Here, we evaluated the genetic overlap between ID and ADHD in an attempt to identify novel ADHD candidate genes. We investigated whether genes affected by rare mutations in ID patients also contribute to ADHD risk through common genetic variation. For this, we used the latest data freeze from the Psychiatric Genomics Consortium (PGC; n=19,210) for discovery and the iPSYCH sample (n=37,076) for replication. To provide functional support for the newly identified ADHD candidates, we used *Drosophila melanogaster*, a model that can facilitate characterization of the involved neural substrates. The role of dopaminergic neurotransmission is well-established in ADHD(1). In addition, circadian genes and circuits have been implicated, as ADHD often goes together with sleep disturbances, and abnormal circadian rhythms of melatonin secretion have been observed in children and adult patients with ADHD(7). Importantly, positive genetic correlations between insomnia and sleep-related traits and ADHD exist(8). Moreover, disrupting the activity of the circadian clock gene *Per1* in both mice and zebrafish revealed ADHD-like symptoms(9). In *Drosophila*, Per1 mutants were deficient for experience-dependent increases in sleep(10). We therefore set out to investigate potential dopaminergic and circadian rhythm components of the identified phenotypes in *Drosophila*. Dissecting the role of neuronal circuits can help to pinpoint the neurotransmitter systems contributing to ADHD as a first step towards an individualization of treatment. Upon downregulation of gene expression pan-neuronally and in relevant neuronal subsets, we assessed locomotor activity, sleep and related parameters as behavioral readouts, which we have previously established to be relevant for ADHD(11).

## MATERIAL AND METHODS

### Ethics statement

The current study used summary statistics of GWAS meta-analyses (GWAS-MA) that had been approved by the local ethics committees and had the required informed consents, as described in the earlier publications(2, 12).

### Cohorts

The Psychiatric Genomics Consortium (PGC) ADHD GWAS meta-analysis (GWAS-MA) data, which were used at the discovery stage in this study, were available as autosome-wide summary statistics, including single nucleotide polymorphism (SNP) data with corresponding P-values and odds ratios. Data were based on nine studies including 5,621 cases and 13,589 controls. Samples were of Caucasian or Han Chinese origin and contained patients meeting ADHD-diagnostic criteria according to the DSM-IV (**Supplementary Table 1**). Detailed procedures of DNA isolation, whole-genome genotyping, and imputation have been described previously(13). Shortly, genome-wide data were obtained from different genotyping arrays (**Supplementary Table 1**) and was imputed using 1000 Genomes Project Phase 3 as a reference panel (NCBI build 37 (hg19) coordinates) for autosomal SNPs. Meta-analytic data were processed through a stringent quality control pipeline applied at the PGC(13).

The gene-set association was replicated in an independent cohort from the Lundbeck Foundation Initiative for Integrative Psychiatric Research (iPSYCH) - Statens Serum Institut (SSI) – Broad ADHD working group (n=37,076)(2).

A meta-analysis of the two data-sets described above (20,183 cases and 35,191 controls) has recently been published as part of the ADHD Working Group of the PGC and the ADHD iPSYCH-SSI-Broad collaboration(2) (https://www.med.unc.edu/pgc/results-and-downloads). This meta-analytic data-set was used by us to perform a gene-based look-up of three genes of interest, using MAGMA software, as described below. Detailed quality control and imputation parameters have been described in the original publication(2). In short, summary data only included markers with a quality (INFO score) >0.8, minor allele frequency (MAF) >0.01, and supported by an effective sample size greater than 70% (8,047,420 markers)(2).

### GWAS of ADHD symptom scores in the Nijmegen Biomedical Study

From the Nijmegen Biomedical Study, a population-based survey in adults(14), data on hyperactivity/impulsivity and inattention symptoms from the self-report DSM-IV-based ADHD-RS(15) and whole-genome genotyping was available for >2,978 individuals. (**Supplementary Table 4**). Detailed information on the sample, procedures of DNA isolation, whole-genome genotyping, and imputation are described in the **Supplementary Methods**. Genome-wide association analysis was performed using a linear regression under an additive model in PLINK v1.9(16, 17) using Ricopili (https://sites.google.com/a/broadinstitute.org/ricopili/). Age, gender, and ten principal components were included as covariates. For subsequent gene-based analyses, SNPs with an INFO score ≥0.8 and MAF ≥0.01 were included.

### ID gene selection

For the selection of the ID gene-set, we used the publicly available ‘Intellectual Disability Gene Panel’ of the Radboudumc department of Human Genetics’ Genome Diagnostics division (downloaded from https://issuu.com/radboudumc/docs/ngs-intellectual_disability_panel_1?e=28355229/50899368 on March 27^th^, 2014). This gene panel listed 490 ID-related genes (shown in **Supplementary Table 2**), based on findings of *de novo* mutations in patients with ID visiting the Radboudumc, collaborating institutes and on literature/public databases. This list forms the basis for diagnostic testing using exome sequencing at Radboudumc.

### Gene-based and gene-set analysis

Genome-wide summary statistics of ADHD (PGC and iPSYCH ADHD GWAS-MA) were used as input for gene-based analyses. We used two software packages to test whether the ID gene-set was associated with ADHD risk. Firstly, the Hybrid set-based test (HYST) of the Knowledge-based mining system for Genome-wide Genetic studies (KGG) version 3.5 software(18) was used for association testing (**Supplementary Methods**). Secondly, the Multi-marker Analysis of GenoMic Annotation (MAGMA) software version 1.02(19) was used (**Supplementary Methods**). The analyses were carried out in two steps. In step 1, the combined effect of the SNPs in (the vicinity of) all ID genes was analyzed. Post hoc, in step 2, the potential effects of the individual genes were investigated, by reviewing their gene-based test-statistics. Genes were considered gene-wide significant if they reached the Bonferroni correction threshold adjusted for the number of genes tested (P<0.000128).

### Functional characterization of *MEF2C* and *TRAPPC9* in *Drosophila melanogaster*

#### Drosophila strains and breeding

*Drosophila* orthologues were retrieved from NCBI protein DELTA-BLAST and ENSEMBL gene-tree (20, 21). The *Drosophila* orthologues of *MEF2C* (termed *Mef2*) and *TRAPPC9* (termed *brun*) were targeted by RNA interference (RNAi)-mediated knockdown using the *UAS-GAL4* system. Tissue-/cell type-specific knockdown was achieved using tissue-/cell type-specific promoters driving GAL4-expression. Several neuronal populations were targeted: *nSyb-GAL4* (*yw*\* *UAS-Dcr-2 hs*(X);; *nSyb-GAL4*)(22) targeting all neurons (pan-neuronal driver), *tim-GAL4* (; *tim*-GAL4, *UAS-Dcr-2*/CyO;)(23) targeting *timeless*-expressing cells including circadian neurons, and *ple-GAL4* (w*; *UAS-Dcr-2*; *ple-GAL4*), obtained from Bloomington *Drosophila* Stock Center #8848, targeting tyrosine hydroxylase-expressing (dopaminergic) neurons, visualized in **Supplementary Fig. 1**. A copy of *UAS-Dcr-2* was incorporated to improve knockdown efficiency(22). The driver stocks were crossed with *UAS*-RNAi lines obtained from the Vienna *Drosophila* Resource Center: v12482 (w^1118^; *UAS-dTRAPPC9*^RNAi^;), v15549 (w^1118^;; *UAS-dMef2*^RNAi-1^), v15550 (w^1118^;; *UAS-dMef2*^RNAi-2^), and v60000 (w^1118^). Progeny of the latter crosses served as genetic background controls. Non-induced UAS-RNAi lines were generated by crossing UAS-RNAi stocks with the isogenic line iso^31^ (Bloomington stock #5905: *w^1118^*;;), replacing the driver in the cross. The driver lines expression pattern was validated by driving GFP expression and UAS-RNAi lines were by qPCR (**Supplementary method**, **Supplementary Fig. 1**). All flies were maintained on standard corn meal food at 28°C with 60% relative humidity in a 12-hour light:dark cycle.

#### Locomotor activity monitoring and calculation of activity and sleep parameters

Locomotor activity of individual 3–5 days old male flies was recorded with the *Drosophila* Activity Monitoring system (Trikinetics, Waltham, USA). The flies were collected with the aid of CO2 and allowed to recover for 24 hours. The activity count was recorded for four days at 28°C and 60% RH in 12-hour light:dark cycle, followed by two days in constant darkness. The activity data was collected every 30-seconds and analyzed in 1-minute bins. Activity and sleep were analyzed with pySolo software(24), defining sleep as ≥5 minutes of inactivity. The average daily activity and sleep were then plotted in 10- and 30-minute bins, respectively. pySolo software was modified to analyze the total activity and sleep between 180–540 min Zeitgeber Time (ZT) for the relative day and 900–1260 min ZT for the relative night to capture periods of stable activity and sleep, as described previously(11). Activity while awake, sleep bout counts, sleep bout duration, and sleep latency were extracted using Sleep and Circadian Analysis Matlab Program (SCAMP)((25)). Data of individual flies from at least two independent experiments were pooled and t-tests were performed with Welch’s correction, when variances were unequal. Results were considered significant if they reached the Bonferroni correction threshold adjusted for the number of drivers tested (P<0.0167). To compare the relative day and night activity, the delta (Δ) activity and sleep between knockdown and genetic background control were calculated (Δ^day^=knockdown^day^–control^day^ and Δ^night^=knockdown^night^–control^night^). All statistical analyses were performed with GraphPad 5.03 Software (San Diego, CA).

## RESULTS

### Association of ID gene-set with ADHD risk

To select candidate genes for the ID gene-set, we used the publicly available ‘Intellectual Disability Gene Panel’. Genes were included based on findings of *de novo* mutations in patients with ID visiting the Radboudumc and collaborating institutes and on literature/public databases (n=490; **Supplementary Table 2**). The set of ID genes was tested for association with ADHD using two different software algorithms in a discovery-replication design. For discovery, we used the PGC ADHD genome-wide association study meta-analysis (GWAS-MA) data (n=19,210) and the KGG software; the HYST test revealed that the ID gene-set as a whole was significantly associated with ADHD risk (P_KGG_=0.0001; n_genes_=387). To assess the robustness of our findings, we tested the association of the ID gene-set with ADHD in the PGC data using the MAGMA software. The results also showed a significant association of the ID gene-set with ADHD risk in the self-contained test (P_self-contained_=0.0412; n_genes_=392), but not in the competitive test (P_competitive_=0.9522). As an independent replication, we tested the gene-set association in the iPSYCH cohort (n=37,076) using MAGMA; the results robustly replicated the significance in the self-contained test (P_self-contained_=1.2429×10^−13^; n_genes_=393). The competitive test was negative again (P_competitive_=0.5306).

To identify the major contributors (i.e. most significantly associated individual genes) to the observed association for further validation, we performed individual gene-wide testing within the gene-set using the PGC data. The most consistent findings across algorithms were for the Myocyte Enhancer Factor 2C gene (*MEF2C*; P_KGG_=1.3×10^−5^ and P_MAGMA_=1.497×10^−4^; **Fig. 1A**), the Trafficking Protein Particle Complex 9 gene (*TRAPPC9*; P_KGG_=7.81×10^−7^ and P_MAGMA_=0.0035; **Fig. 1B**), and the ST3 Beta-Galactoside Alpha-2,3-Sialyltransferase 3 gene (*ST3GAL3*; P_KGG_=6.18×10^−5^ and P_MAGMA_=6.808×10^−4^; **Fig. 1C**). Genebased p-values for all genes in both KGG and MAGMA analyses can be found in **Supplementary Table 3**. A look-up in the recently published combined PGC+iPSYCH GWAS-MA(2) revealed genome-wide significant results for gene-wide analysis of *ST3GAL3* and *MEF2C*, and nominal significance for *TRAPPC9* (P_*ST3GAL3*_=4.6406×10^−13^, P_*MEF2C*_=2.671×10^−10^, and P_*TRAPPC9*_=0.0184).

**Figure 1:**
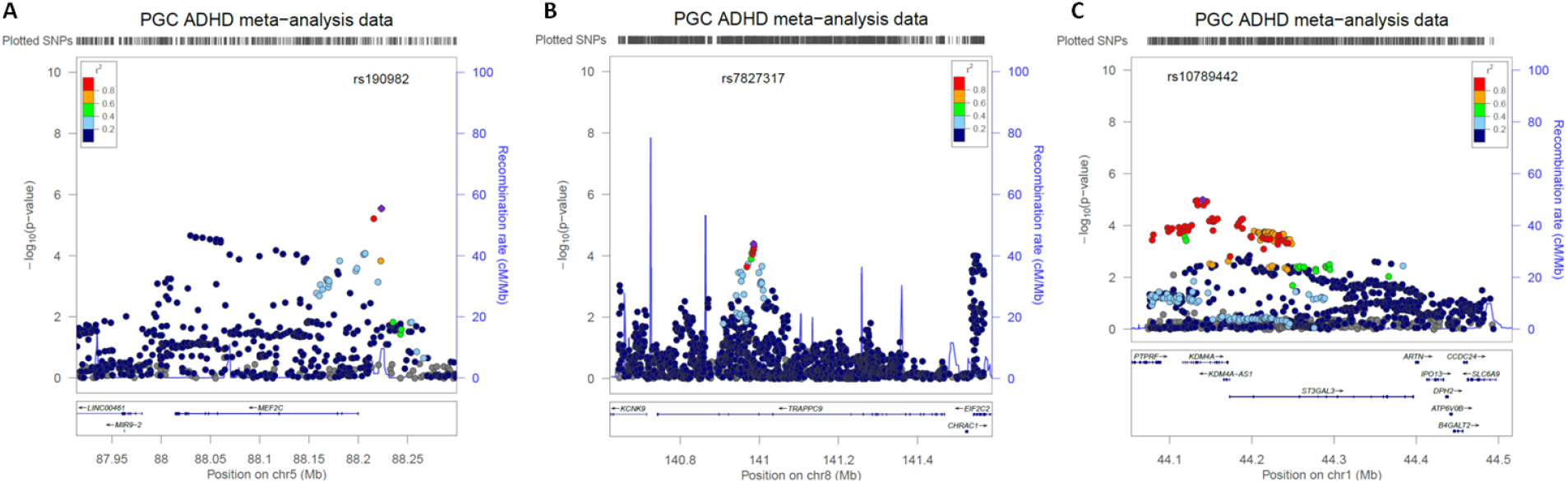
Regional association plots showing association signals for ADHD in the PGC GWAS-MA (n=19,210) for the three most consistently associated genes, including flanking regions of 100 kb. (**A**) *MEF2C* locus with the top-SNP (rs190982) indicated by the purple dot. (**B**) *TRAPPC9* locus with the top-SNP (rs7827317) indicated by the purple dot. (**C**) *ST3GAL3* locus with the top-SNP (rs10789442) indicated by the purple dot. Results are shown as −log (p-value) for genotyped and imputed SNPs. The color of each marker reflects its LD (r^2^) with the strongest associated SNP (in purple). The recombination rate is plotted in blue. cM/Mb, centimorgan/megabase. Chr, chromosome.

To distinguish between contributions of the two separate symptom domains of ADHD, hyperactivity/impulsivity and inattention, we investigated gene-based associations in a population-based cohort of > 2,978 adults(14). *MEF2C* and *TRAPPC9* showed gene-based association with hyperactive/impulsive ADHD symptoms, but not inattentive ADHD symptoms. However, these results did not survive correction for multiple testing (**Supplementary Table 5**).

### Functional validation and characterization of MEF2C and TRAPPC9 in Drosophila

Next, we investigated the validity of the newly identified ADHD candidate genes by mapping their effects on ADHD-related phenotypes in the fruit fly *Drosophila melanogaster*. As mutations in these gene are already proven to cause monogenic forms of intellectual disability but not ADHD, we focused our efforts on the ADHD-related phenotypes. We did so by investigating neuronal subsets, in addition to pan-neuronal knockdown of the genes. This allowed us to characterize the different circuits through which individual ADHD risk genes may act. Secondly, it may reveal phenotypes that might otherwise be masked by opposing actions of different neurons in the same circuit(26). We have earlier established *Drosophila* as a model for ADHD by showing that pan-neuronal knockdown of ADHD genes preferentially caused (dopamine-related) increased locomotor activity and sleep loss at night(11). Two of the three ADHD candidate genes were found conserved in *Drosophila*: the *MEF2* gene-family homolog *Mef2* (further referred to as *dMEF2*) and the *TRAPPC9* homolog *brun* (further referred to as *dTRAPPC9*). *ST3GAL3* is found in vertebrates, and no known orthologue has been identified in *Drosophila*. We investigated locomotor activity and sleep after knocking down *dMEF2* and *dTRAPPC9* expression in all neurons, or more specifically in dopaminergic or circadian neurons. Cell type-specific knockdown was achieved by driving the expression of RNA interference (RNAi) in the neuronal populations of interest (pan-neuronal, dopaminergic, and circadian) using the binary UAS/Gal4 system. Flies were monitored in 12-hour light:dark scheme, mimicking day and night period. We also investigated behavior in 24-hour constant darkness conditions, given our earlier model that the dopamine-related increased locomotor activity is present in the absence of light(11).

### dMEF2 knockdown gives rise to elevated night-time activity and sleep defects

Pan-neuronal knockdown of *dMEF2* expression caused no changes in activity and sleep during the stable period of the relative day compared to the genetic background control (**Fig. 2A**), but significantly increased night activity (P_activity_=0.0059) and reduced sleep (P_sleep_=0.014) (**Fig. 2A, Supplementary Table 6**). In constant darkness, the knockdown also showed significantly increased activity (P_activity_=0.0088) and less sleep (P_sleep_=0.00037) in the relative night period (**Fig. 2B, Supplementary Table 7**). This increased activity was the result of increased activity counts per waking minute (**Fig. 2A, B**). Further analysis of sleep parameters revealed a tendency of reduced sleep bout duration and an increased sleep latency in the relative night period (**Supplementary Fig. 3A, B**). Knockdown of *dMEF2* in dopaminergic neurons showed increased night activity (P_activity_=1.8×10^−15^) and reduced sleep (P_sleep_=5.1×10^−15^; **Fig. 2C, Supplementary Table 6**). Activity and sleep during the relative day period were not different from the genetic background control (**Fig. 2C**). In constant darkness, increased activity and reduced sleep were observed in both relative day (P_activity_=4.5×10^−7^, P_sleep_=2.6×10^−17^) and night (P_activity_=1.6×10^−17^, P_sleep_=9.5×10^−26^) (**Fig. 2D**, **Supplementary Table 7**). Under these conditions, the increased activity was exclusively driven by a sleep defect; activity while awake was even lower than in controls (**Fig. 2C, D**). The sleep defect was accompanied with an increased sleep bout count and a reduced sleep bout duration (**Supplementary Fig. 3C, D**). Knockdown using *tim-GAL4* did not yield viable flies, precluding further analysis. The increased activity and sleep loss were the result of induced knockdown of the gene of interest as non-induced UAS-RNAi lines showed no increased activity or sleep defect (**Supplementary Fig. 5**).

**Figure 2:**
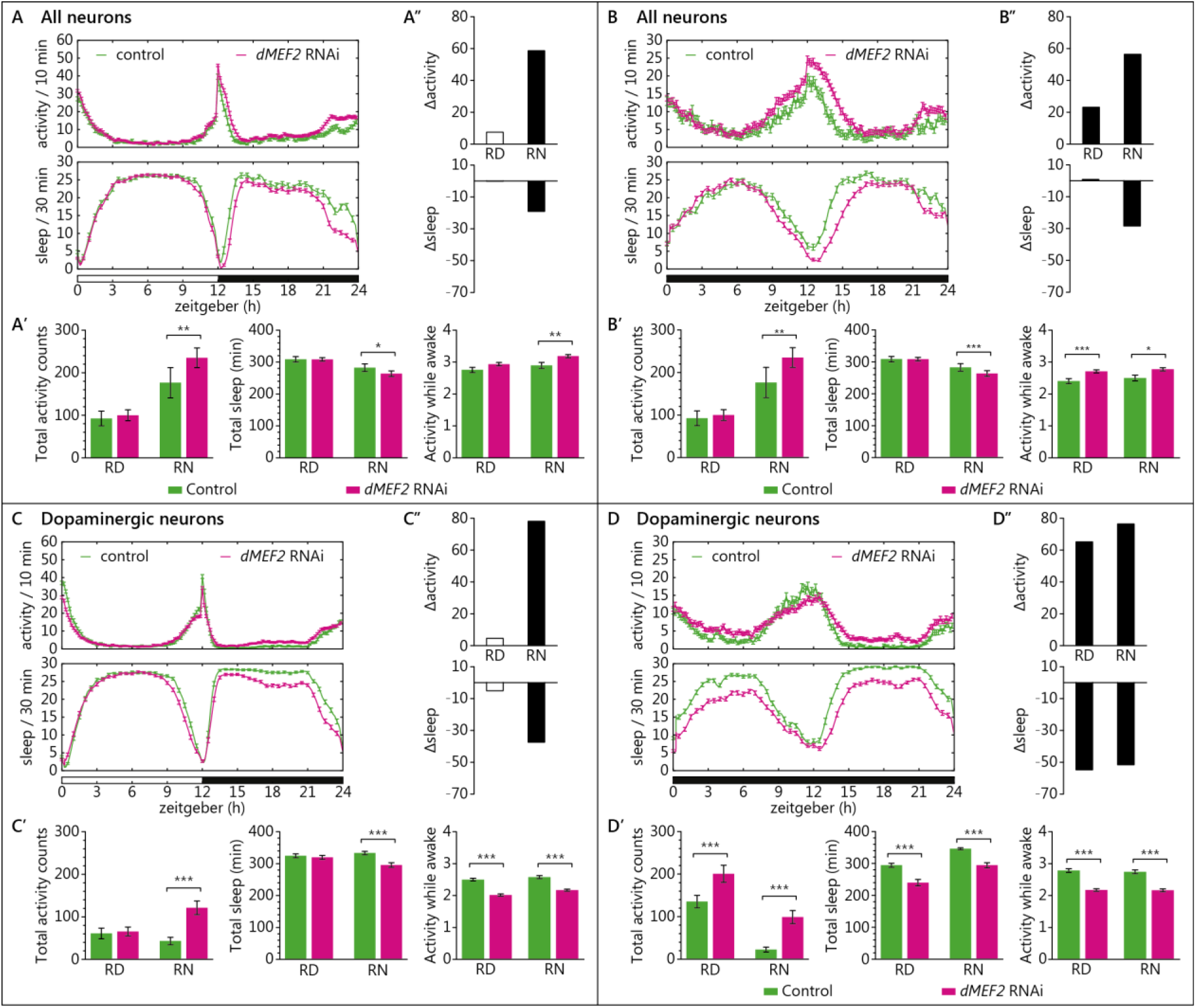
Knockdown of *dMEF2* in all neurons, or in dopaminergic neurons specifically, results in higher activity and reduced sleep in the relative night. **(A, B)** Activity and sleep plot of pan-neuronal *dMEF2* knockdown, **(A)** in 12-hour light:dark cycle and **(B)** in constant darkness. **(A’, B’)** Quantification of total activity, sleepand activity while awake during stable periods (RD: zeitgeber 3-9h, RN: zeitgeber 15-21h), excluding the activity peaks (zeitgeber 0-3h, 9-15h and 21-24h). Panneuronal knockdown of *dMEF2* showed increased activity, activity while awake and sleep loss during the RN period during both 12-hour light:dark cycle and constant darkness. **(A”, B”)** Δ_activity_ and Δ_sleep_: the findings for 12-hour light:dark cycle and for constant darkness both reveal that the difference between groups is greater in the absence of light. **(C, D)** Activity and sleep plot of dopaminergic neuron *dMEF2* knockdown, **(C)** in 12-hour light:dark cycle and **(D)** in constant darkness. **(C’, D’)** Quantification of total activity, sleep and activity while awake during stable periods (RD: zeitgeber 3-9h, RN: zeitgeber 15-21h), excluding the activity peaks (zeitgeber 0-3h, 9-15h and 21-24h). Dopaminergic neuronal knockdown of *dMEF2* showed increased activity and sleep loss in the RN period during 12-hour light:dark cycle and both in RD and RN during constant darkness. The knockdown showed lower activity while awake than the control in both RD and RN in 12-hour light:dark cycle and also in constant darkness. **(C”, D”)** Δ_activity_ and Δ_sleep_: the findings for 12-hour light:dark cycle and for constant darkness reveal that the difference is greater when light is absent. For the figure, data from two *dMEF2* lines with identical *UAS*-RNAi constructs were combined since the results from the individual lines are highly consistent; the individual data are presented in **Supplementary Fig. 2**. Further activity and sleep parameters are shown in **Supplementary Fig. 3.** RD, relative day; RN, relative night. Error bars represent standard error of means (SEM). N=3 biological replicates, minimum 20 flies/replicate; *P<0.0167 (Bonferroni correction threshold), **P<0.01, ***P<0.001.

### dTRAPPC9 knockdown influences activity and sleep only in neuronal subtypes

Pan-neuronal knockdown of *dTRAPPC9* did not result in observable alterations in activity or sleep in either the 12-hour light:dark cycle (**Fig. 3A, Supplementary Fig. 3A**) or in constant darkness (**Fig. 3B, Supplementary Fig. 3B**). Specific knockdown of *dTRAPPC9* in dopaminergic neurons caused significantly reduced activity and increased day sleep during the relative day (P_activity_ =0.0022; P_sleep_=0.013), but not in the night (**Fig. 3C, Supplementary Table 6**). In constant darkness, relative night activity was increased and sleep was reduced (P_activity_=0.012; P_sleep_=0.015; **Fig. 3D, Supplementary Table 7**). In contrast, knockdown of *dTRAPPC9 timeless*-expressing neurons resulted in increased night activity and reduced night sleep (P_activity_=4.2×10^−5^; P_sleep_=0.00022; **Fig. 3E, Supplementary Table 6**). In constant darkness, increased activity and reduced sleep were also present in the relative night (P_activity_=0.00017; P_sleep_=0.010; **Fig. 3F, Supplementary Table 7**). This increased activity was the result of higher activity counts per waking minute (**Fig. 3E, F**). Further analysis of sleep parameters revealed an increased sleep bout count and an increased sleep latency in the relative night period (**Supplementary Fig. 4E, F**). The activity and sleep loss were the result of knockdown of the gene of interest as non-induced UAS-RNAi did not show increased activity or sleep defects (**Supplementary Fig. 5**).

**Figure 3:**
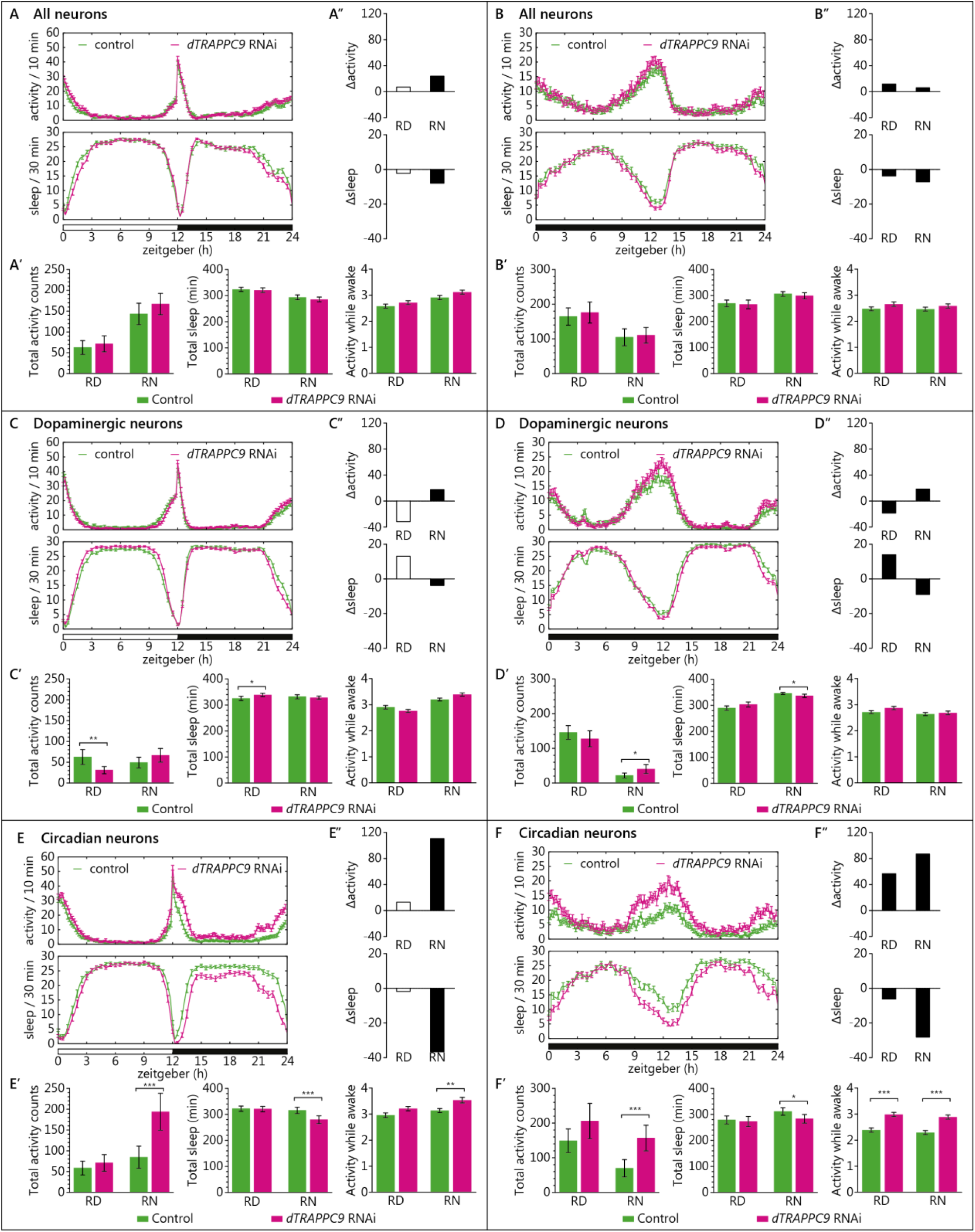
Knockdown of *dTRAPPC9* results in higher activity and reduced sleep, when induced in circadian rhythm neurons, but not in all or dopaminergic neurons. **(A, B)** Activity and sleep plot of pan-neuronal *dTRAPPC9* knockdown, **(A)** in 12-hour light:dark cycle and **(B)** in constant darkness. **(A’, B’)** Quantification of total activity, sleep and activity while awake during stable periods (RD: zeitgeber 3-9h, RN: zeitgeber 15-21h), excluding the activity peaks (zeitgeber 0-3h, 9-15h and 21-24h). Panneuronal knockdown of *dTRAPPC9* showed similar activity, sleep, and activity while awake. **(A”, B”)** Δ_activity_ and Δ_sleep_. **(C, D)** Activity and sleep plot of dopaminergic neuron *dTRAPPC9* knockdown, **(C)** in 12-hour light:dark cycle and **(D)** in constant darkness. **(C’, D’)** Quantification of total activity, sleep and activity while awake during stable periods (RD: zeitgeber 3-9h, RN: zeitgeber 15-21h), excluding the activity peaks (zeitgeber 0-3h, 9-15h and 21-24h). Dopaminergic neuronal knockdown of *dTRAPPC9* showed lower activity and increased sleep in the RD period during 12-hour light:dark cycle. During constant darkness, the knockdown showed higher activity and sleep loss in the RN period. **(C”, D”)** Δ_activity_ and Δ_sleep_. **(E, F)** Activity and sleep plot of circadian rhythm neuron *dTRAPPC9* knockdown, **(E)** in 12-hour light:dark cycle and **(F)** in constant darkness. **(E’, F’)** Quantification of total activity, sleep and activity while awake during stable periods (RD: zeitgeber 3-9h, RN: zeitgeber 15-21h), excluding the activity peaks (zeitgeber 0-3h, 9-15h and 21-24h). Circadian neuronal knockdown of *dTRAPPC9* showed increased activity and sleep loss in the RN period during both 12-hour light:dark cycle and constant darkness. The knockdown showed increased activity while awake in the RN period during 12-hour light:dark cycle and both RD and RN period in constant darkness. **(E”, F”)** Δ_activity_ and Δ_sleep_: the findings for 12-hour light:dark cycle and for constant darkness reveal that the difference is greater when light is absent. Further activity and sleep parameter are shown in **Supplementary Fig. 4**. RD, relative day; RN, relative night. Error bars represent standard error of means (SEM). N=3 biological replicates, minimum 20 flies/replicate; *P<0.0167 (Bonferroni correction threshold), **P<0.01, ***P<0.001.

## DISCUSSION

In the current study, we used a robust discovery-replication design in the currently largest available, independent data sets to show that genes affected by rare genetic variation in ID patients also contribute to ADHD risk through common genetic variation. In the discovery phase, we also used different algorithms to test gene-set association to further test the robustness of findings. In the KGG HYST test and MAGMA, we found significance in both self-contained tests but not the competitive test. A non-significant competitive p-value in the competitive test should be interpreted as an inability to disentangle the part of the polygenicity attributable to the genes in the gene-set from the polygenicity “remaining” (i.e. not captured by the set) on the rest of the genome. In combination with a significance in the self-contained test, it should not be interpreted as no effect of the selected gene-set on the outcome. Our replication in the larger, independent data-set makes this point convincingly. Even more convincing is the fact that two of the three novel ADHD candidate genes that we identified, *MEF2C* and *ST3GAL3*, are among the genome-wide significant findings in the recently published ADHD GWAS-MA(2).

Interestingly, our study design produced reproducible findings in much smaller sample sizes than those needed to reach genome-wide significance, which makes such overlap studies an attractive source of genes which have not previously been implicated by GWAS in ADHD. While we based our selection of ID genes on a diagnostic gene panel, many more ID genes are currently being discovered through the fast advances in next generation sequencing technology; those surely leave additional ADHD genes to be identified.

Our interdisciplinary approach, combining highly powered statistical analyses in humans with functional analyses in an unconventional, validated *Drosophila* model for ADHD-related behavior(11, 27), allowed for a direct validation and further characterization of neural substrates involved. None of our three top-genes had been investigated in the context of ADHD before. *MEF2C* encodes a member of the MADS box transcription factor, which binds to the conserved MADS box sequence motif(28). *MEF2C* is important for normal neuronal function by regulating neuronal proliferation, differentiation, survival, and synapse development(29, 30). It also plays a role in hippocampal-dependent learning and memory, possibly by controlling the number of excitatory synapses(31). While both haplo-insufficiency and gene-duplications of *MEF2C* give rise to ID in humans, most severe ID cases are linked to large deletions removing part or all of *MEF2C* and *de novo* point mutations in the gene(32); individuals with duplications of *MEF2C* usually display a milder phenotype, with only mild cognitive impairment(33). This is why we chose to model reduced gene-expression in *Drosophila* in this study. Common variants (SNPs) in the *MEF2C* locus have previously been found associated with various cognitive, neuropsychiatric, and neurodegenerative phenotypes, such as intelligence(34), schizophrenia(35), and Alzheimer’s disease(36), indicating pleiotropic effects of this gene on a range of phenotypes. The findings of our study add ADHD to this list and suggest that this is linked to the role of *MEF2C* in neurotransmission contributing to it through dopaminergic neurons. However, knowing that in *Drosophila dMEF2* expression is important in maintaining normal circadian rhythm(37, 38), we cannot yet rule out an additional role of *dMEF2* in circadian neurons in the ADHD-related behaviors, as our *dMEF2* knockdown did not yield flies.

*TRAPPC9* has been implicated in NF-kB signaling and is possibly involved in intracellular trafficking. *TRAPPC9* is highly expressed in postmitotic neurons of the cerebral cortex, and MRI analysis of affected patients showed defects in axonal connectivity(39). The *Drosophila TRAPPC9* has been studied for its involvement in meiotic division in *Drosophila* male gametes(40), but a neuronal function has not been described so far. *TRAPPC9*-associated ID is linked to loss of function of the gene(41). Hyperactive behavior has so far been reported in one patient with a *TRAPPC9* mutation(42). Our findings indicate that *TRAPPC9* can play a role in ADHD and suggest that the gene primarily acts by affecting neurons involved in circadian regulation. Interestingly, while the *dTRAPPC9* dopaminergic neuron knockdown showed similar night activity and sleep profile to the control, somewhat lower activity and increased sleep was observed during the day, suggesting *dTRAPPC9* having several, cell-type specific roles in the brain.

While both *dMEF2* and *dTRAPPC9* pan-neuronal knockdown showed weak or no activity and sleep phenotype compared to the control, the knockdown in specific neuronal populations did result in pronounced alterations in activity and sleep profiles. This is consistent with earlier work reporting that tissue-specific knockdown can lead to more severe outcomes compared to null mutants(43). Importantly, Sitaraman and coworkers(26) previously identified distinct neuronal subtypes within the part of the *Drosophila* brain that oppositely regulates sleep. Considering that the whole nervous system is a mixture of various neuronal populations, each with specific function in specified phases of development, the different activity and sleep profiles of *dMEF2* and *dTRAPPC9* knockdown in distinct neuronal populations indicates the need to investigate gene function in different brain circuits and identifies a particular strength of our study. Importantly, our findings also show that different behavioral characteristics can contribute to ADHD-like activity phenotypes downstream of different gene defects; the *dMEF2* knockdown showed increased activity and sleep loss as a result of sleep defects, while in *dTRAPPC9* knockdown flies the altered activity and sleep was caused by hyperactivity.

The third ADHD candidate we identified, *ST3GAL3*, is not conserved in *Drosophila*, hence we were not able to study its contribution to ADHD-relevant behavior. The gene encodes a membrane protein (ST3GalIII) that adds sialic acid to the terminal site of glycolipids or glycoproteins. The gene is expressed in a variety of tissues including neurons(44). In mice, *St3gal2* and *St3gal3* are responsible for nearly all the terminal sialyation of brain gangliosides and play an important role in cognition(44). A role in brain development is also likely in humans, as the human brain is particularly enriched in sialic acid-containing glycolipids (i.e. gangliosides)(45). Gangliosides are known to modulate calcium homeostasis and signal transduction in neurons(46). Common genetic variants in *ST3GAL3* have also been associated with educational attainment(47). Interestingly, in a recent study of DNA-methylation, sites annotated to *ST3GAL3* were found associated with ADHD symptom trajectories in the population (48). The use of alternative animal models, e.g. mouse or zebrafish, is warranted to characterize the neuronal circuits underlying *ST3GAL3*’s effects on ADHD-related behavior.

In the current study, we modeled one of the two behavioral symptom domains of ADHD, namely hyperactivity. This was consistent with our findings – though only nominally significant, likely due to limited sample size – that *MEF2C* and *TRAPPC9* were more strongly associated with hyperactivity/impulsivity than with inattention. However, being able to assess gene effects related to the second domain, i.e. inattention, would likely help to elucidate additional neural substrates and circuits involved in ADHD. Currently, there are multiple paradigms to assess attention available in *Drosophila*, as summarized by de Bivort and van Swinderen(49).

In summary, the genetic overlap we observed between ID and ADHD may suggest biological pleiotropy, in which genetic variation severity in an overlapping set of genes is linked to the severity of neurodevelopmental phenotypes. Functional characterization of neural substrates involved revealed that the novel ADHD candidate genes may impact disease etiology through different biological pathways.

## Disclosures and acknowledgments

### Acknowledgements

Funding: Part of this work was carried out on the Dutch national e-infrastructure with the support of SURF Foundation. The Nijmegen Biomedical Study is a population-based survey conducted at the Department for Health Evidence and the Department of Laboratory Medicine of the Radboud University Medical Center. Principal investigators of the Nijmegen Biomedical Study are LALM Kiemeney, ALM Verbeek, DW Swinkels and B Franke. The authors would like to acknowledge grants supporting their work from the Netherlands Organization for Scientific Research (NWO), i.e. the NWO Brain & Cognition Excellence Program (grant 433-09-229), the TOP grant Program (grant 912-12-109 to AS), the Veni Innovation Program (grant 91-614-084 to MV), and the Vici Innovation Program (grant 016-130-669 to BF). Additional support is received from the Dutch National Science Agenda for the NWANeurolabNL project (grant 400 17 602), from the European Community’s Seventh Framework Programme (FP7/2007 – 2013) under grant agreement n° 602805 (Aggressotype), from the European Community’s Horizon 2020 Programme (H2020/2014 – 2020) under grant agreements n° 643051 (MiND), n° 667302 (CoCA), and n° 728018 (Eat2beNICE), by the ECNP Network ADHD across the Lifespan, and from the Lundbeck Foundation (grant numbers R102-A9118 and R155-2014-1724) to iPSYCH. Computational resources for handling and statistical analysis of iPSYCH data on the GenomeDK HPC facility were provided by the Centre for Integrative Sequencing, iSEQ, Aarhus University, Denmark (grant to ADB). The ADHD working group of the Psychiatric Genomics Consortium (PGC) and the iPSYCH-SSI-Broad collaboration ADHD Working Group contributed with independent sets of summary statistics of consortium findings. The data and a complete list of contributing samples and people can be obtained from the PGC download website (https://www.med.unc.edu/pgc/results-and-downloads).

## Author contributions

Study conception and supervision: M.K., E.S., A.S., M.vdV., B.F. Obtained funding: A.D.B., L.A.K., A.S., M.vdV., B.F. Provided samples and/or data: D.D., A.D.B., L.A.K., A.CN. Conducted analyses: M.K., E.S., A.vR., D.D., N.R.M. Contributed to analyses and data interpretation: H.G.B, A.A.-V., L.A.K Writing group: M.K., E.S., M.vdV., B.F. All authors reviewed and approved the final version of this manuscript.

