## Supplementary Material for "Intellectual Disability-related genes increase ADHD risk and locomotor activity in *Drosophila*"

[$] shared final responsibility

**The supplementary information contains:**

- Supplementary methods
- Supplementary Figures 1 to 5
- Supplementary Table 1 to 7
- References

### SUPPLEMENTARY METHODS

#### GWAS of ADHD symptom scores in the Nijmegen Biomedical Study

The Nijmegen Biomedical Study is a population-based survey conducted at the Department of Health Evidence, and the Department of Laboratory Medicine of the Radboud university medical center(1). Within the NBS cohort information using the ADHD-RS(2), a self-report DSM-IV-based symptom list with high validity in population-based and case samples, was collected. (**Supplementary Table 4**). Total ADHD hyperactivity/impulsivity and inattention symptoms were transformed, by using the Blom-transformation, to achieve a normal distribution. Detailed procedures of DNA isolation have been described previously(1). Genome-wide genotyping was carried out using the Illumina HumanOmniExpress-12 platform. Quality control, imputation and association analyses were done using the bioinformatics pipeline Ricopili (https://sites.google.com/a/broadinstitute.org/ricopili/). Stringent quality control was applied to the cohort following standard procedures for GWAS, including filters for call rate, Hardy-Weinberg equilibrium and heterozygosity rates. The NBS cohort was then phased and imputed using the 1000 Genomes Project phase 3 (1KGP3)(3, 4) imputation reference panel using Eagle(5) and Minimac3(6), respectively. Cryptic relatedness and population structure were evaluated using a set of high-quality markers pruned for linkage disequilibrium (LD). Genetic relatedness was estimated using PLINK v1.9(7, 8) to identify first and second-degree relatives and one individual was excluded from each pair. Genetic outliers were identified for exclusion based on principal component analysis using EIGENSOFT (version 5.0.1)(1, 9, 10). Genome-wide association analysis has been performed using a linear regression under an additive model in PLINK v1.9 (7, 8) using Ricopili (https://sites.google.com/a/broadinstitute.org/ricopili/). Age and gender were included as covariates. The first ten principal components were added to account for possible population stratification effects. For subsequent gene-based analyses, only SNPs with an INFO score ≥0.8 and MAF ≥0.01 were included.

**Gene-based and gene-set analysis**

We extracted SNPs and association p-values from the PGC ADHD GWAS-MA for the ID gene-set. Since this meta-analysis only covered the autosomes, X-chromosomal genes were excluded, leaving 396 autosomal genes. All SNPs lying within these genes (according to UCSC hg19 position(11)), including flanking regions of 100 kb to capture regulatory regions, were extracted. In total, 308,952 SNPs with a MAF ≥0.01 and INFO-score ≥0.8 were considered for further analysis. The gene-based and gene-set association analyses in the iPSYCH data were performed at secured servers in Denmark at the GenomeDK high performance-computing cluster (http://genome.au.dk) by the responsible researcher using the same protocols.

We used two software packages to test whether the ID gene-set was associated with ADHD risk. Firstly, the Hybrid set-based test (HYST) of the Knowledge-based mining system for Genome-wide Genetic studies (KGG) version 3.5 software(12) was used for association testing. Within this software package, we chose the Hybrid set-based test (HYST)(13) for association testing. A text file listing all 396 autosomal ID genes and a text file listing all SNPs that were extracted from the PGC ADHD GWAS-MA, were used as input for KGG. Imputed and quality-controlled genome-wide genotyping data of the Brain Imaging Genetics (BIG) cohort(14) were used as a reference to define the underlying linkage disequilibrium (LD) structure. The LD upper limit was set to a $r^2$ of 0.8, while the lower limit was set to 0.2. Next, a gene-based association scan was run with HYST, based on a hybrid test of Gates and a scaled Chi-square test, in which SNPs without LD information were ignored. Gene-set and gene-based p-values were calculated for 388 genes, since eight genes could either not be annotated by the reference file (*ATP1A2, ERCC2, GRIK2, GSS, KIAA1279, MMACHC*, and *TCF4*) or had too few SNPs left after quality control (*B4GALT7*).

Secondly, the Multi-marker Analysis of GenoMic Annotation (MAGMA) software version 1.02(15) was used. First, genome-wide SNP data from a reference panel (1000 Genomes, v3 phase1(3)) were annotated to NCBI Build 37.3 gene locations using a symmetric 100 kb flanking window, and both files were downloaded from http://ctglab.nl/software/magma. Next, the gene annotation file was used to map the genome-wide SNP data to assign the SNPs to the genes and to calculate gene-based p-values for each cohort separately. For the gene-based analyses, single SNP p-values within a gene were transformed into a gene-statistic by taking the mean of the $\chi 2$-statistic among the SNPs in each gene. To account for LD, the 1000 Genomes Project European sample was used as a reference to estimate the LD between SNPs within (the vicinity of) the genes (http://ctglab.nl/software/MAGMA/ref_data/g1000_ceu.zip). Gene-wide p-values were converted to z-values reflecting the strength of the association of each gene with the phenotype, with higher z-values corresponding to stronger associations. Subsequently, we tested, whether all 396 ID-related genes in the gene-set are jointly associated with each of the phenotypes, using an intercept-only linear regression model including a subvector corresponding to the genes in the gene-set. This self-contained analysis evaluated, whether the regression coefficient of this regression was larger than 0, testing whether the gene-set contains any association at all. To test if the genes in the gene-set are more strongly associated with each phenotype than other genes, the regression model was then expanded including all genes outside the gene-set. This competitive test tested, whether the association of a gene-set is different from association of genes outside the gene-set. To account for the potentially confounding factors of gene size and gene density, both gene size and gene density as well as their logarithms were included as covariates in the competitive gene-set analysis. Four genes were not included in the analyses, because they were either missing from the annotation file (*CHKB-CPT1B* and *SOX2-OT*) or contained too few SNPs (*B4GALT7* and *RAB3GAP2*).

The analyses were carried out in two steps. In step 1, the combined effect of the SNPs in (the vicinity of) all ID genes was analyzed. Post hoc, in step 2, the potential effects of the individual genes were investigated, by reviewing their gene-based test-statistics. Genes were considered gene-wide significant if they reached the Bonferroni correction threshold adjusted for the number of genes tested ($P<0.000128$).

### Validation of *Drosophila* models

#### *Drosophila* Immunolabellings

The driver line expression pattern was validated by using the driver to drive GFP expression in the adult *Drosophila* brain. Female virgins of the driver lines were crossed with the males of a membrane-GFP expression line (Bloomington stock #5137: $y^1w^*$; *UAS-mCD8::GFP.L*). Flies were grown at 25°C, 70% relative humidity (RH). Brains of 3–5 day-old males were dissected, rinsed with phosphate buffered saline (PBS), and fixed in 3.7% paraformaldehyde at room temperature for 30 minutes. The fixed brains were rinsed with 0.3% Triton X-100 in PBS (PBST) and then blocked in blocking buffer containing 3% normal goat serum (NGS) in PBST at room temperature for 1 hour. Brains were then incubated with primary antibody rabbit anti-GFP (Invitrogen) in blocking buffer (1:600) at 4°C for 72h and washed five times in PBST at room temperature for 10 minutes. Subsequently, brains were incubated with secondary antibody Alexa488 goat anti-rabbit (Molecular Probes) in blocking buffer (1:700) at 4°C for 48h and washed with PBST at room temperature for 5x 10 minutes, before being mounted on slides with mounting medium ProLong Gold Antifade (Life Technologies). Images of the brains were taken in stacks at 200x magnification with Zeiss Axio Imager microscope (with Apotome.2); exposure time was kept constant. The maximum projections of the images were created using FIJI 1.51n(16).

#### mRNA extraction and Quantitative Real Time PCR (qRT-PCR)

The efficiency of the utilized *dMEF2* RNAi lines was previously validated (17-20). The efficiency of the *dTRAPPC9* RNAi line had not previously been established. Therefore, we validated the *dTRAPPC9* RNAi efficacy by driving *dTRAPPC9* knock down with the ubiquitous driver (Bloomington stock #4414: $w^*$ *UAS-*

*Dcr2*; *Act5C-Gal4*/CyOGFP; ) and measuring the difference of the *dTRAPPC9* transcript level. The males of the RNAi and genetic background lines were crossed with the virgins of the ubiquitous driver line, raised at 25°C, 70% RH. For qRT-PCR, 3–5 days old male progeny were collected with $CO_2$, snap-frozen, and the total RNA was isolated with RNeasy Lipid Tissue kit (Qiagen). The isolated RNA concentration was quantified with Nanodrop (Thermo Fisher Scientific). A total of 200 ng RNA was converted to cDNA with the iScript kit (BioRad). The cDNA was used as a template for qRT-PCR using the SybrGreen kit (Applied Biosystems) in 96-well plate format in a 7900HT Fast Real-Time PCR system (Applied Biosystems). The following primers were used in qRT-PCR: *β'COP* (forward: 5'-aactacaacaccctggagaagg-3'; reverse: 5'- acatcttctcccaattccaaag-3'), *RpII215* (forward: 5'- ccgcgatacttctctccac-3'; reverse: 5'-gaccagctaggcgacattc-3'), *dTRAPPC9* (forward: 5'- aaagtgcgacgtgtcaatcac-3'; reverse: 5'-gtgctccacaggatgatcgt-3'). A threshold was carefully applied to determine the cycle threshold ($C_T$) values using SDS 2.4 software (Applied Biosystems). The fold change of dTRAPPC9 was calculated with ΔΔCt method(21). Significance was calculated using paired T-test in GraphPad 5.03 (San Diego, CA).

**URLs**

https://www.med.unc.edu/pgc/results-and-downloads

http://ctglab.nl/software/MAGMA/ref_data/g1000_ceu.zip

http://locuszoom.sph.umich.edu/

https://issuu.com/radboudumc/docs/ngs-intellectual_disability_panel_1?e=28355229/50899368

**DATA AVAILABILITY**

GWAS summary statistics used in the paper are available directly from the web for ADHD GWAS-MA data of the PGC ADHD Working Group and the combined ADHD GWAS-MA data of the PGC ADHD Working Group of the PGC + ADHD iPSYCH-SSI-Broad collaboration (https://www.med.unc.edu/pgc/results-and-downloads). The ID gene-set can be downloaded from https://issuu.com/radboudumc/docs/ngs-intellectual_disability_panel_1?e=28355229/ 50899368 on March 27th, 2014. All data needed to evaluate the conclusions in the paper are present in the paper or the supplementary materials.

**SUPPLEMENTARY FIGURES**

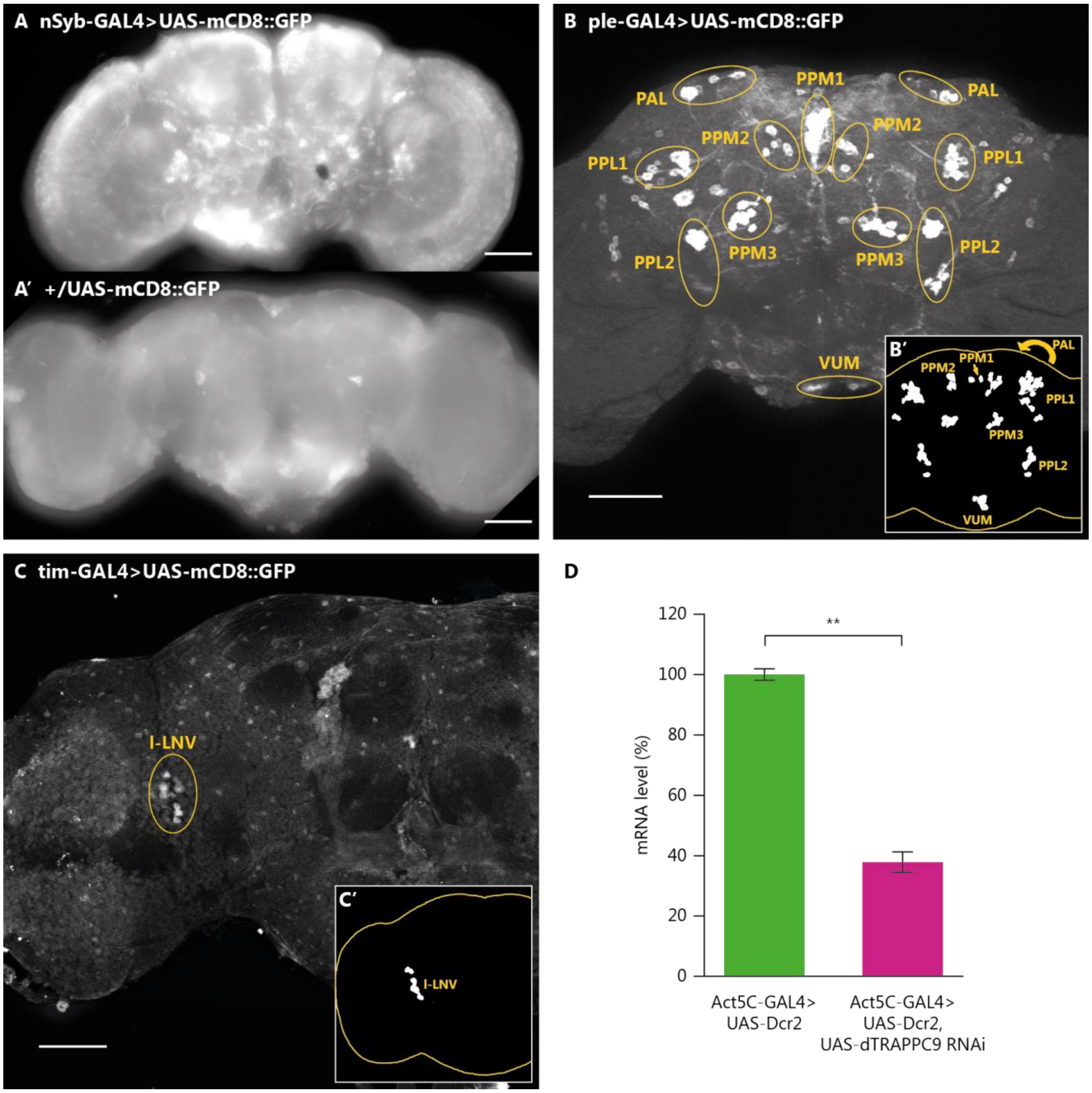

**Supplementary Figure 1.** Validation of the *Drosophila* lines used in this study. **(A)** Maximum projection of adult male brains of membrane GFP expression driven by *nSyb-GAL4* (*nSyb-GAL4>UAS-mCD8::GFP)*, and of **(A')** non-induced *UAS-mCD8::GFP (+/UAS-mCD8::GFP)*, illustrating background staining. Using same exposure time, the intensity of fluorescent signal from *nSyb-GAL4>UAS-mCD8::GFP* is higher compared to non-induced *UAS-mCD8::GFP*; the broad expression of *nSyb* can be appreciated as labelling of granular appearance throughout most parts of the brain. Maximum projection of adult male brains of membrane GFP expression driven by **(B)** *ple-GAL4* (*ple-GAL4>UAS-mCD8::GFP*) and **(C)** *tim-GAL4* (*tim-*

*GAL4>UAS-mCD8::GFP*). Sketches of (B') *ple-GAL4* and (C') *tim-GAL4* expression pattern, illustrating their expression pattern with identifyable neurons, matching the patter described in the literatures(22, 23). (**A-C**) The fluorescent patterns of all three drivers, *nSyb-Gal4*, *ple-GAL4* and *tim-GAL4* matches their described expression pattern. All scale bars indicate 50 µm. **(D)** Relative expression of dTRAPPC9 mRNA in the indicated genotypes (driver control in green, *dTRAPPC9* knock down in magenta), as determined by quantitative real-time PCR (qRT-PCR). Both values have been normalized against the control condition. 3 days old adult male were used; Ubiquitous knockdown of *dTRAPPC9* (*Act5C-GAL4> UAS-Dcr2*, *UAS-dTRAPPC9*$^{RNAi}$) results in reduced expression of *dTRAPPC9* to 40% compared to control (*Act5C-GAL4>UAS-Dcr2*). n=3 biological replicate; error bars represent standard error of means (SEM); **p<0.01.

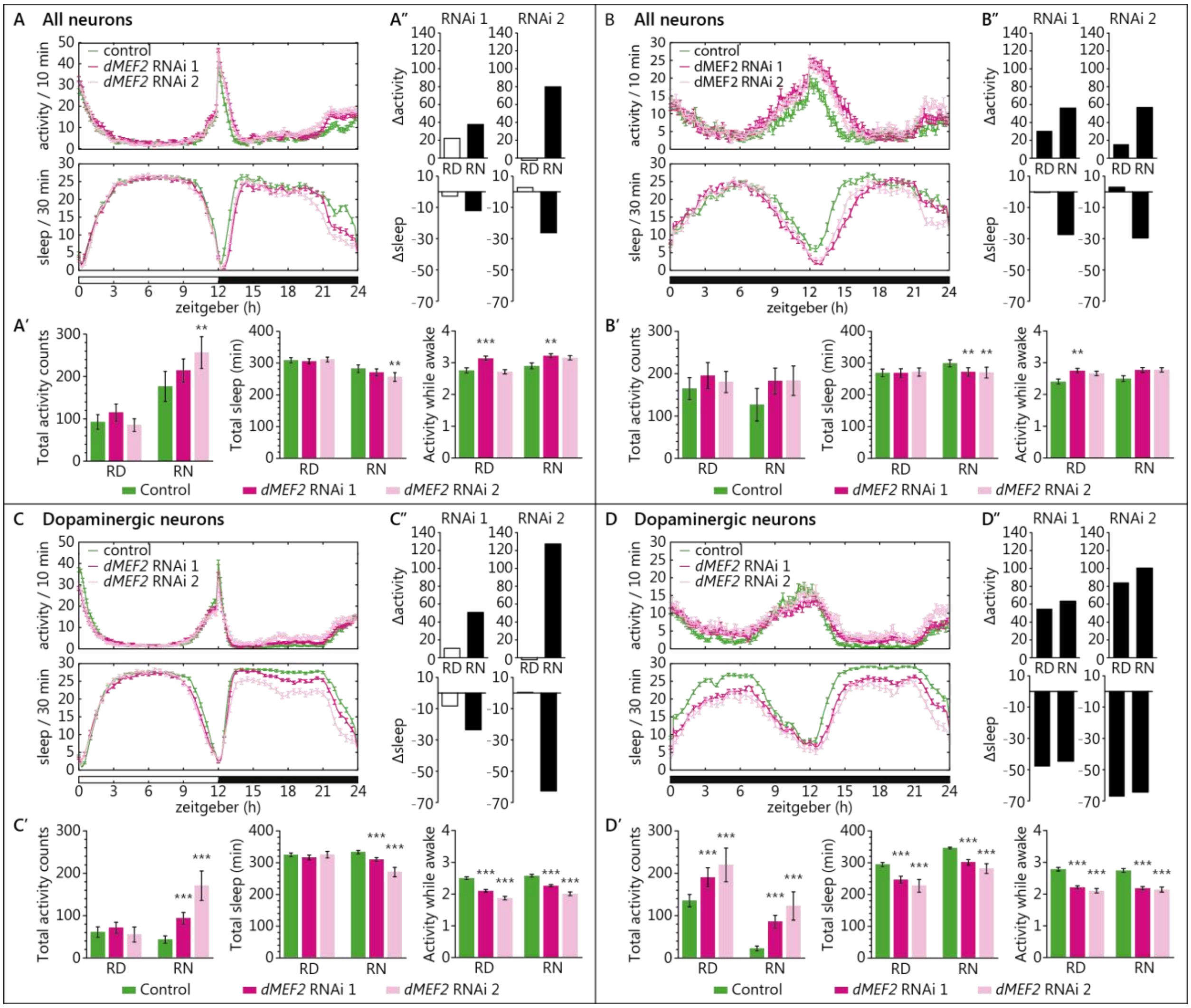

**Supplementary Figure 2**: Knockdown of *dMEF2* in all neurons using two independent RNAi lines (A, B), or specifically in dopaminergic neurons (C, D), resulted in higher activity and reduced sleep in the relative night. **(A, B)** Activity and sleep plots of pan-neuronal *dMEF2* knockdown, **(A)** in 12-hour light:dark cycle and **(B)** constant darkness. **(A′, B′)** Quantification of total activity, sleep, and activity while awake during stable periods (RD: zeitgeber 3-9h, RN: zeitgeber 15-21h) excluding the activity peaks (zeitgeber 0-3h, 9-15h, and 21-24h). Pan-neuronal knockdown of *dMEF2* with both RNAi lines showed a tendency towards increased activity and activity while awake, as well as sleep loss during the RN period in both 12-hour light:dark cycle and constant darkness. **(A″, B″)** $\Delta_{activity}$ and $\Delta_{sleep}$: the findings for 12-hour light:dark cycle and for constant darkness both reveal that the difference between groups is greater in the absence of light. **(C, D)** Activity and sleep plot of dopaminergic neuron *dMEF2* knockdown, **(C)** in 12-hour light:dark cycle and **(D)** constant darkness. **(C', D')** Quantification of total activity, sleep and activity while awake during stable periods (RD: zeitgeber 3-9h, RN: zeitgeber 15-21h) excluding the activity peaks (zeitgeber 0-3h, 9-15h and 21-24h). Dopaminergic neuronal knockdown of *dMEF2* with both RNAi lines showed increased activity and decreased sleep in the RN period and both in RD and RN during constant darkness. The knockdown showed lower activity while awake than the control in both RD and RN in 12-hour light:dark cycle and also in constant darkness. **(C", D")** $\Delta_{activity}$ and $\Delta_{sleep}$: the findings for 12-hour light:dark cycle and for constant darkness both reveal that the difference between groups is greater in the absence of light. RD, relative day; RN relative night. Error bars represent standard error of means (SEM). N=3 biological replicates, minimum 20 flies/replicate; *P<0.0167 (Bonferroni correction threshold), **P<0.01, ***P<0.001.

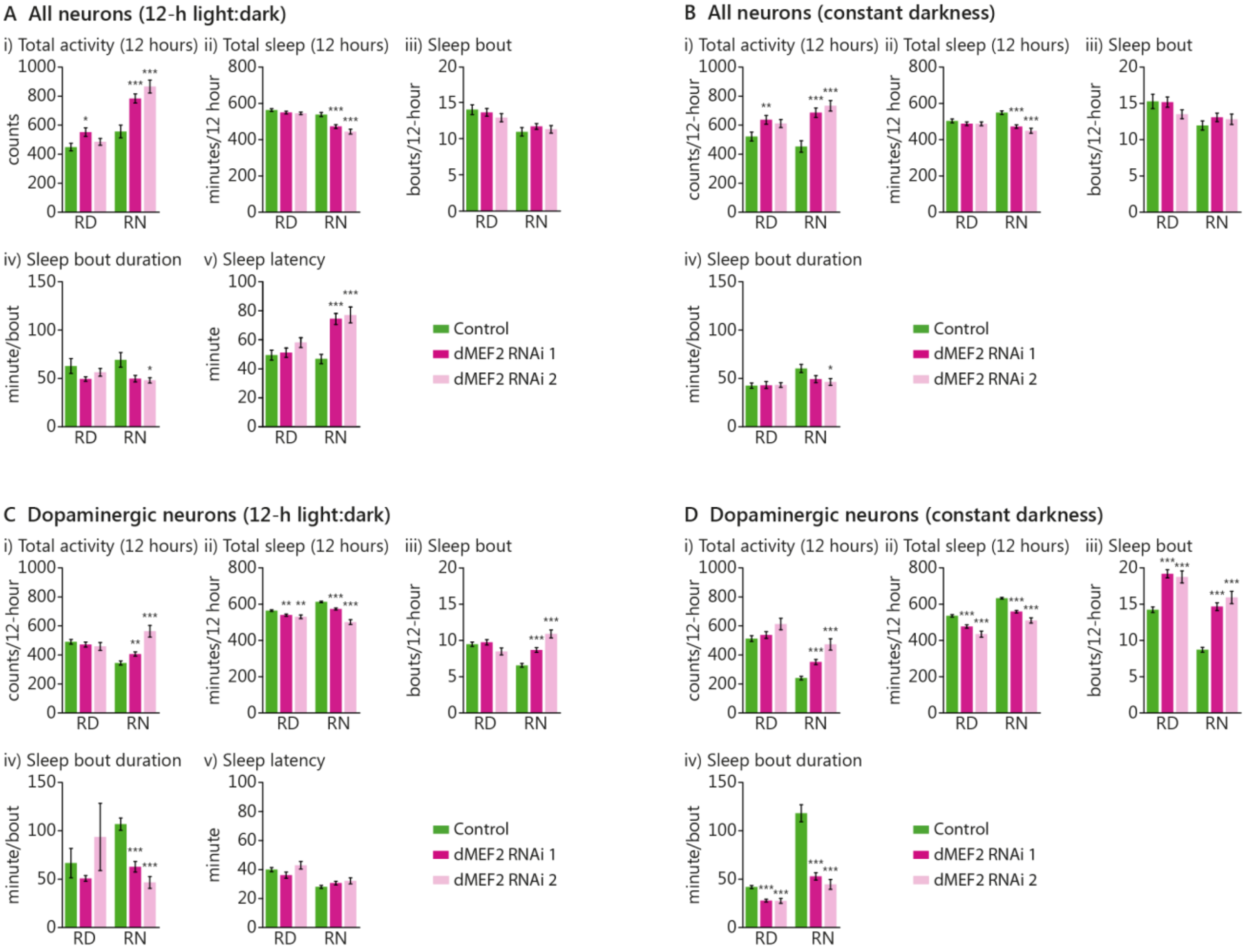

**Supplementary Figure 3**. Quantification of activity and sleep parameters of *dMEF2* knockdown, calculated over the entire 12-hour day and 12-hour night. **(A,B)** Activity and sleep parameters of pan-neuronal knockdown of *dMEF2* monitored in **(A)** 12-hour light:dark cycle and **(B)** constant darkness. **(C,D)** Activity and sleep parameters of dopaminergic neuron knockdown of *dMEF2* monitored in **(C)** 12-hour light:dark cycle and **(D)** constant darkness. Sleep latency parameter is only available for 12-h light:dark cycle. RD, relative day; RN, relative night. Error bars represent standard error of means (SEM) *P<0.0167 (Bonferroni correction threshold), **P<0.01, ***P<0.001.

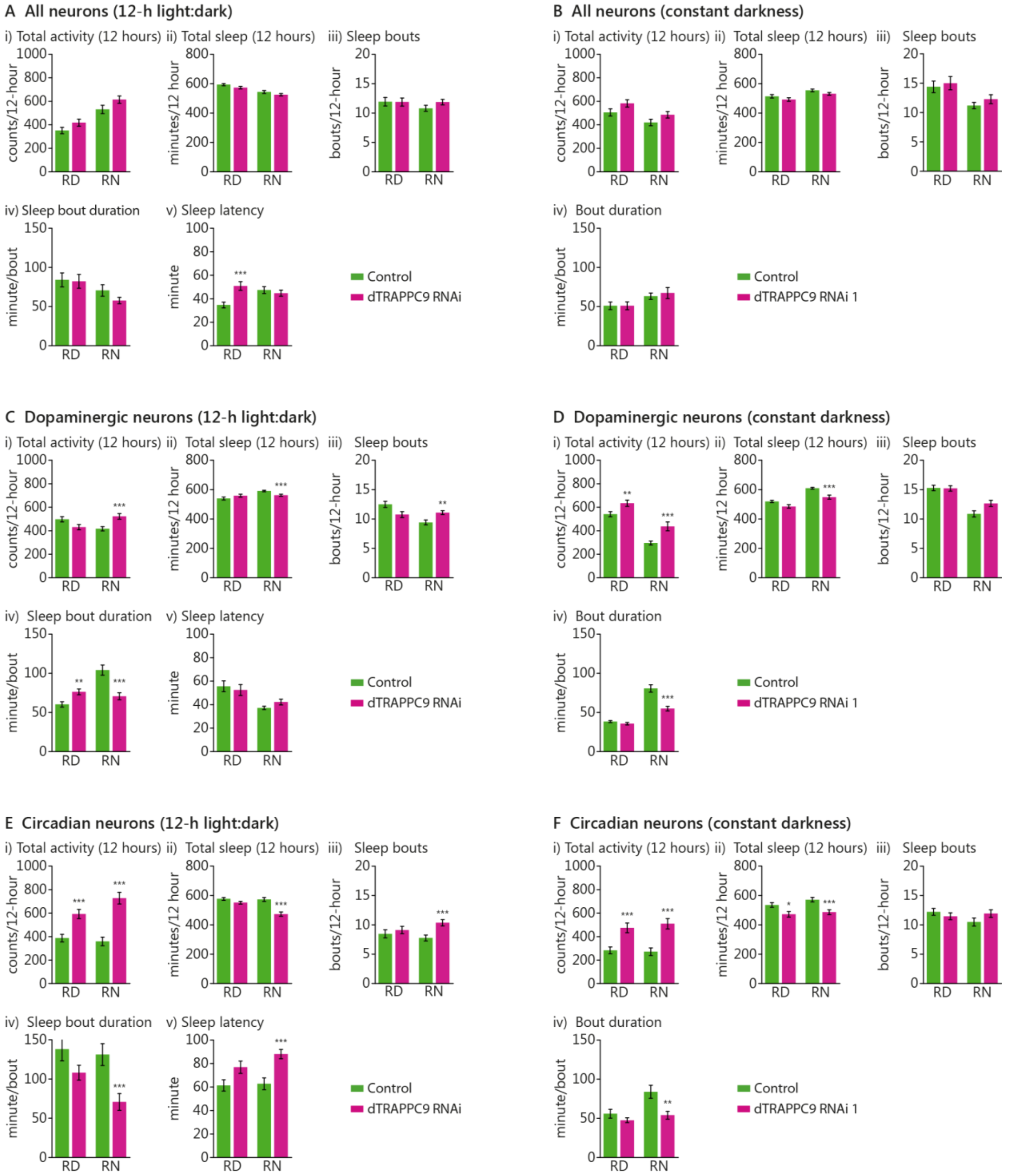

**Supplementary Figure 4.** Quantification of activity and sleep parameters of *dTRAPPC9* knockdown, calculated for the entire 12-hour day and 12-hour night. **(A,B)** Activity and sleep parameters of pan-neuronal knockdown of *dTRAPPC9* monitored in **(A)** 12-hour light:dark cycle and **(B)** constant darkness.

**(C,D)** Activity and sleep parameters of dopaminergic neuron knockdown of *dTRAPPC9* monitored in **(C)** 12-hour light:dark cycle and **(D)** constant darkness. **(E,F)** Activity and sleep parameters of circadian neuron knockdown of *dTRAPPC9* monitored in **(E)** 12-hour light:dark cycle and **(F)** constant darkness. Sleep latency parameter is only available for 12-h light:dark cycle. RD, relative day; RN, relative night. Error bars represent standard error of means (SEM) *P<0.0167 (Bonferroni correction threshold), **P<0.01, ***P<0.001.

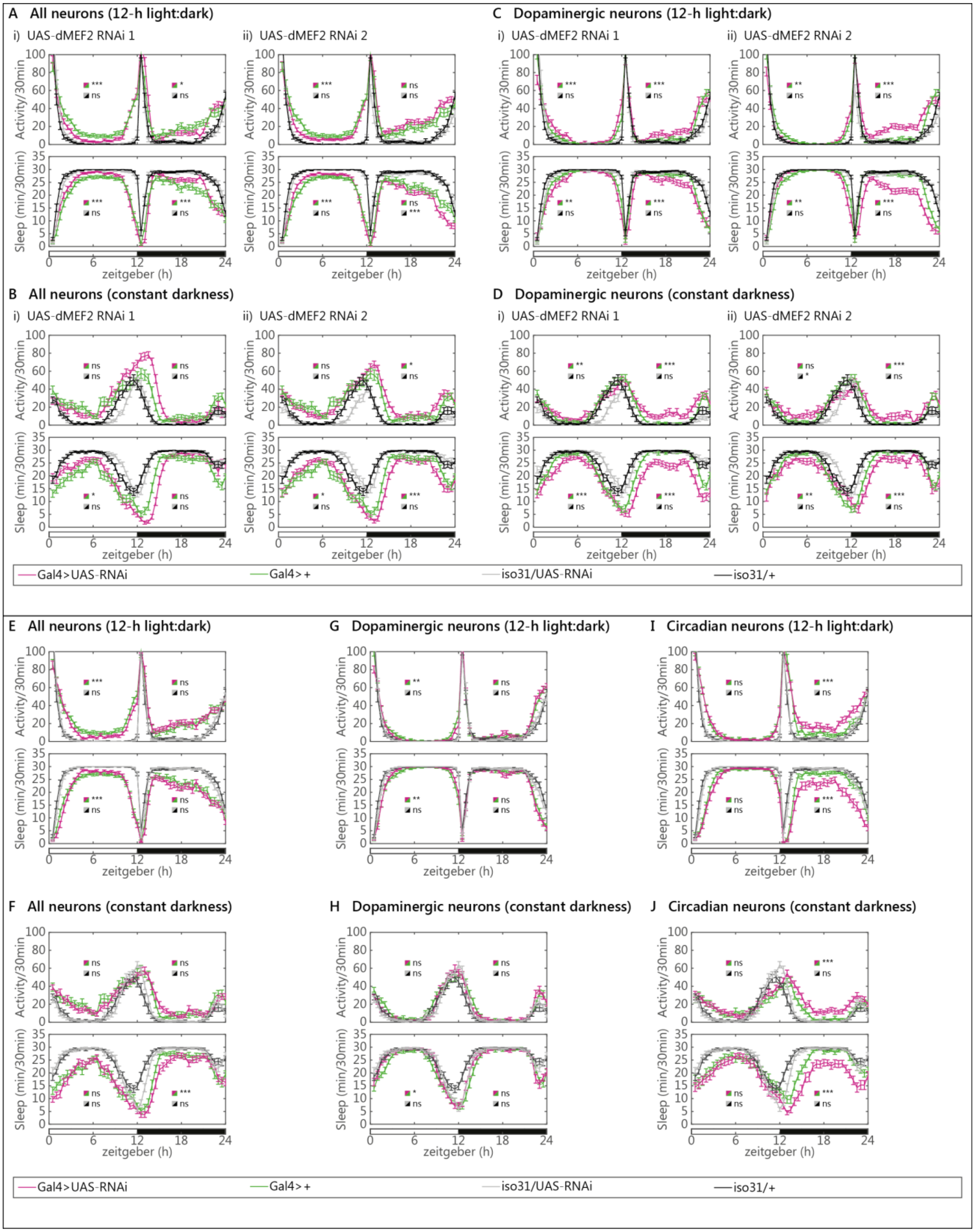

**Supplementary Figure 5.** Uninduced *dMEF2* and *dTRAPPC9* RNAi lines have similar activity and sleep to the genetic background control, demonstrating specificity of obtained phenotypes. **(A, B, C, D)** Activity and sleep profile of *dMEF2* knockdown in **(A,B)** all neurons and **(C,D)** dopaminergic neurons. **(E,F,G,H,I,J)** Activity and sleep profile of dTRAPPC9 knockdown in **(E,F)** all neurons and **(G,H)** dopaminergic neurons and **(I,J)** circadian neurons, monitored in **(A,C,E,G,I)** 12-h light:dark scheme and **(B,D,F,H,J)** constant darkness.

**SUPPLEMENTARY TABLES**

**Supplementary Table 1**. Description of individual PGC ADHD cohorts as included in the PGC ADHD GWAS-MA.

| **Study name** | | **Study design** | **Ancestry** | **Cases (N[a])** | **Controls (N[a])** | **Genotyping platform** | **Reference[b]** |
|---|---|---|---|---|---|---|---|
| Short | Full sample name | | | | | | |
| CARD | UK sample | Case/control | Caucasian | 727 | 5,081 | Illumina Human660W-Quad BeadChip (ADHD cases) and Illumina Human 1.2M BeadChip (controls) | (24) |
| CHIN | Chinese sample | Case/control | Han Chinese | 1,040 | 963 | Affymetrix 6.0 array | (25) |
| CHOP | Children's Hospital of Philadelphia | Trio/control | European descent | 335 trios | 2026 | Illumina Infinium II HumanHap550 BeadChip | (26) |
| GERM | ADHD patient sample consisted of children and adolescents Aachen, Cologne, Essen, Marburg, Regensburg, and Würzburg | Case/control | Caucasian (all German) | 495 | 1,300 | Illumina Human660W-Quadv1(ADHD cases) and Illumina HumanHap550v3 (controls) | (27) |
| CROS | Toronto University, SickKids project | Trio | Mainly Caucasian | 170 | | Affymetrix 6.0 array | (28) |
| IMAGE1 | Phase I of the International Multisite ADHD Genetics | Trio | Western European | 909 trios | | Perlegen Sciences 600 K | (29) |

| | | | | | | | |
|---|---|---|---|---|---|---|---|
| | Project | | origin | | | | |
| IMAGE2 | Phase II of IMAGE | Case/control | Predominantly European origin | 896 | 2,455 | Affymetrix 5.0 array (ADHD cases) and Affymetrix 6.0 arrays (controls) | (30) |
| PuWMa | Pfizer funded study from the University of California, Los Angeles, Washington University and the Massachusetts General Hospital | Trio | Mainly Caucasian | 735 trios | | Illumina Human1M or Human1M-Duo BeadChip platform | (31) |
| SPAN | Spanish sample | Case/control | Caucasian | 607 | 584 | Illumina HumanOmni1-Quad BeadChip platform | (32) |
| **TOTAL** | | | | **5,621** | **13,589** | | |

[a] Based on sample of primary publication [b] Primary publication reporting individual study sample.

**Supplementary Table 2**. ID gene panel listing 490 candidate genes for ID

*>>> see file supplementary_table_S2.xls <<<*

**Supplementary Table 3**. Results of gene-based association analyses for 396 ID-related genes with ADHD risk in the PGC ADHD GWAS-MA data (n=19,210). Results for both KGG and MAGMA analyses, using a 100 kb flanking region, are shown.

*>>> see file supplementary_table_S3.xls <<<*

N/A: no gene-based result obtained for this gene.

**Supplementary Table 4**. Descriptive information for the Nijmegen biomedical Study (NBS) cohort and for phenotype assessment (1).

| N (% F) | Age (SD) | Mean HI score (SD) | Mean IA score (SD)* |
|---|---|---|---|
| 3019 (53.7%) | 58.7695 (16.29241) | 4.6706 (3.02432) | 4.4495 (3.41122) |

HI=Hyperactivity/impulsivity symptoms, IA=inattention symptoms, * N=2,978

**Supplementary Table 5**. Results of gene-based association analyses with hyperactivity/impulsivity in the NBS cohort (N=3,019 for HI analyses and N=2,978 for IA analyses)

| Gene | P Hyperactivity/impulsivity | P Inattention |
|---|---|---|
| *MEF2C* | 0.016852 | 0.79178 |
| *TRAPPC9* | 0.022311 | 0.18314 |
| *ST3GAL3* | 0.69772 | 0.46721 |

None of the gene-based association results remains significant after correction for multiple testing (P<0.00833).

**Supplementary Table 6.** Summary of results of functional characterization of *dMef2* and *dTRAPPC9* knockdown in *Drosophila* in 12-hour light:dark cycle.

| | | Activity | | | | Sleep | | | | Activity while awake | | | |
|---|---|---|---|---|---|---|---|---|---|---|---|---|---|
| | Number of flies (N) | Total RD | P | Total RN | P | Total RD | P | Total RN | P | Total RD | P | Total RN | P |
| **Pan-neuronal** | | | | | | | | | | | | | |
| ***dMef2* control** | 62 | 92.63 | | 176.65 | | 308.69 | | 282.78 | | 2.76 | | 2.899043 | |
| ***dMef2* RNAi** | 135 | 100.13 | 0.50 | 235.25 | 0.0059 | 308.66 | 1.0 | 263.72 | 0.014 | 2.94 | 0.0552 | 3.187696 | 0.0066* |
| ***dMef2* RNAi 1** | 68 | 114.76 | 0.10 | 214.30 | 0.092 | 306.92 | 0.64 | 270.71 | 0.14 | 3.14 | 0.0004 | 3.218168 | 0.0059* |
| ***dMef2* RNAi 2** | 67 | 85.29 | 0.52 | 256.51 | 0.0026 | 311.45 | 0.63 | 256.62 | 0.0049 | 2.71 | 0.6869 | 3.154226 | 0.0279* |
| ***dTRAPPC9* control** | 66 | 62.78 | | 143.33 | | 324.20 | | 293.48 | | 2.58 | | 2.911718 | |
| ***dTRAPPC9* RNAi** | 68 | 69.82 | 0.58 | 167.14 | 0.19 | 321.97 | 0.70 | 285.50 | 0.22 | 2.71 | 0.2250 | 3.121813 | 0.0660 |
| **Dopaminergic neurons** | | | | | | | | | | | | | |
| ***dMef2* control** | 132 | 61.09 | | 43.36 | | 324.85 | | 333.36 | | 2.50 | | 2.57905 | |
| ***dMef2* RNAi** | 185 | 65.74 | 0.57 | 121.43 | $1.8x10^{-15}$* | 319.76 | 0.24 | 295.92 | $5.1x10^{-15}$* | 2.02 | $3.11x10^{-16}$* | 2.170843 | $4.26x10^{-12}$* |
| ***dMef2* RNAi 1** | 119 | 71.52 | 0.25 | 94.09 | $3.5x10^{-09}$* | 316.65 | 0.086 | 309.90 | $5.7x10^{-08}$ | 2.10 | $2.25x10^{-10}$* | 2.266093 | $1.43x10^{--7}$* |
| ***dMef2* RNAi 2** | 66 | 55.33 | 0.60 | 170.72 | $8.1x10^{-10}$* | 325.37 | 0.93 | 270.72 | $8.6x10^{-12}$* | 1.87 | $2.07x10^{-14}$ | 2.00965 | $1.66x10^{-11}$ |
| ***dTRAPPC9* control** | 84 | 62.54 | | 49.22 | | 325.63 | | 331.70 | | 2.90 | | 3.19955 | |
| ***dTRAPPC9* RNAi** | 79 | 31.06 | 0.0022* | 66.75 | 0.0931 | 338.74 | 0.013* | 327.93 | 0.47 | 2.76 | 0.1183 | 3.391288 | 0.0233 |
| **Circadian neurons** | | | | | | | | | | | | | |
| ***dMef2* control** | *Not tested* | | | | | | | | | | | | |
| ***dMef2* RNAi** | *Not tested* | | | | | | | | | | | | |
| ***dMef2* RNAi 1** | *Not tested* | | | | | | | | | | | | |
| ***dMef2* RNAi 2** | *Not tested* | | | | | | | | | | | | |

| | | | | | | | | | | | | | |
|---|---|---|---|---|---|---|---|---|---|---|---|---|---|
| ***dTRAPPC9* control** | 56 | 58.30 | | 83.22 | | 322.48 | | 316.39 | | 2.97 | | 3.137899 | |
| ***dTRAPPC9* RNAi** | 53 | 71.22 | 0.19 | 193.96 | 4.2x10-05 | 320.73 | 0.80 | 279.80 | 0.00022 | 3.21 | 0.0405 | 3.536499 | 0.0029* |

*Welch correction was performed. RD=relative day. RN=relative night.

**Supplementary Table 7.** Summary of results of functional characterization of *dMef2* and *dTRAPPC9* knockdown in *Drosophila* in constant darkness.

| | | **Activity** | | | | **Sleep** | | | | **Activity while awake** | | | |
|---|---|---|---|---|---|---|---|---|---|---|---|---|---|
| | Number of flies (N) | Total RD | P | Total RN | P | Total RD | P | Total RN | P | Total RD | P | Total RN | P |
| **Pan-neuronal** | | | | | | | | | | | | | |
| ***dMef2* control** | 62 | 164.99 | | 126.95 | | 269.14 | | 299.56 | | 2.41 | | 2.501068 | |
| ***dMef2* RNAi** | 133 | 188.18 | 0.17 | 183.37 | 0.0088 | 270.45 | 0.95 | 271.17 | 0.00037* | 2.71 | 0.0009 | 2.773957 | 0.01001* |
| ***dMef2* RNAi 1** | 67 | 195.66 | 0.13 | 183.03 | 0.023 | 268.76 | 0.97 | 272.25 | 0.0022 | 2.75 | 0.0012 | 2.773551 | 0.0214 |
| ***dMef2* RNAi 2** | 66 | 180.58 | 0.39 | 183.72 | 0.030 | 272.16 | 0.74 | 270.07 | 0.0047* | 2.66 | 0.0169 | 2.774404 | 0.0169* |
| ***dTRAPPC9* control** | 66 | 164.47 | | 104.77 | | 269.70 | | 306.25 | | 2.48 | | 2.469804 | |
| ***dTRAPPC9* RNAi** | 66 | 176.08 | 0.56 | 110.72 | 0.72 | 269.99 | 0.72* | 299.15 | 0.35* | 2.66 | 0.1155 | 2.588258 | 0.2754 |
| **Dopaminergic neurons** | | | | | | | | | | | | | |
| ***dMef2* control** | 128 | 135.52 | | 22.63 | | 294.56 | | 346.07 | | 2.78 | | 2.744557 | |
| ***dMef2* RNAi** | 173 | 200.79 | 4.5x10$^{-07}$* | 99.08 | 1.6x10$^{-17}$* | 239.79 | 2.6x10$^{-17}$* | 294.47 | 9.5x10$^{-26}$* | 2.18 | 2.04x10$^{-16}$* | 2.169431 | 9.88x10$^{-14}$* |
| ***dMef2* RNAi 1** | 112 | 190.39 | 8.5x10$^{-05}$* | 86.06 | 5.5x10$^{-13}$* | 246.57 | 1.0x10$^{-11}$* | 301.43 | 8.7x10$^{-17}$* | 2.22 | 3.14x10$^{-13}$* | 2.186388 | 9x10$^{-12}$* |
| ***dMef2* RNAi 2** | 61 | 219.88 | 0.00016* | 122.98 | 1.4x10$^{-07}$* | 227.34 | 8.9x10$^{-09}$* | 281.69 | 2.1x10$^{-11}$* | 2.11 | 5.93x10$^{-11}$ | 2.140733 | 2.27x10$^{-8}$ |
| ***dTRAPPC9* control** | 83 | 145.91 | | 21.78 | | 289.57 | | 345.81 | | 2.76 | | 2.641114 | |
| ***dTRAPPC9* RNAi** | 78 | 127.61 | 0.23 | 40.35 | 0.012* | 303.54 | 0.030* | 336.82 | 0.015* | 2.87 | 0.0695 | 2.688091 | 0.6042 |
| **Circadian neurons** | | | | | | | | | | | | | |
| ***dMef2* control** | *Not tested* | | | | | | | | | | | | |
| ***dMef2* RNAi** | *Not tested* | | | | | | | | | | | | |
| ***dMef2* RNAi 1** | *Not tested* | | | | | | | | | | | | |

| | | | | | | | | | | | | | |
|---|---|---|---|---|---|---|---|---|---|---|---|---|---|
| ***dMef2* RNAi 2** | *Not tested* | | | | | | | | | | | | |
| ***dTRAPPC9* control** | 55 | 149.22 | | 69.96 | | 278.91 | | 311.48 | | 2.39 | | 2.292069 | |
| ***dTRAPPC9* RNAi** | 53 | 206.00 | 0.067 | 157.03 | 0.00017 | 272.45 | 0.62 | 283.35 | 0.011 | 2.99 | $6.63x10^{-7}$ | 2.888294 | $5.5x10^{-7}$ |

*Welch correction was performed. RD=relative day. RN=relative night.
